# Suppressor of Fused controls the proliferation of postnatal neural stem and precursor cells via a Gli3-dependent mechanism

**DOI:** 10.1101/438705

**Authors:** Hector G. Gomez, Jesse Garcia Castillo, Hirofumi Noguchi, David Aguilar, Samuel J. Pleasure, Odessa R. Yabut

## Abstract

The ventricular-subventricular zone (V-SVZ) of the forebrain is the source of neurogenic stem/precursor cells for adaptive and homeostatic needs throughout the life of most mammals. Here, we report that Suppressor of Fused (SUFU) plays a critical role in the establishment of the V-SVZ at early neonatal stages by controlling the proliferation of distinct subpopulations of stem/precursor cells. Conditional deletion of Sufu in radial glial progenitor cells (RGCs) at E13.5 resulted in a dramatic increase in the proliferation of Sox2+ Type B cells, as well as Gsx2+ ventral forebrain derived transit amplifying precursor cells (TACs). In contrast, we found a significant decrease in Tbr2+ dorsal forebrain derived TACs indicating that innate differences between dorsal and ventral forebrain derived Type B cells influence Sufu function. However, most precursors failed to survive and accumulated in the dorsal V-SVZ, demonstrating that precursors are unable to transition into functional differentiated progenies. These defects were accompanied by reduced Gli3 expression, yet despite reduced Gli3 levels, activation of Sonic hedgehog (Shh) signaling did not occur implying that the Sufu-Gli3 regulatory axis may influence other signaling pathways in the neonatal dorsal V-SVZ. These data suggest that Sufu plays a critical role in controlling Gli3 function in the establishment and survival of functional stem/precursor cells in the postnatal dorsal V-SVZ.

**SUMMARY STATEMENT:** Conditional deletion of Sufu cause dramatic expansion of neural stem/precursor cells in the neonatal ventricular-subventricular (SVZ) zone. This defect occurs through a Gli3-dependent mechanism that results in the downregulation of Shh signaling.

## INTRODUCTION

Tissue-specific stem cell niches persist at postnatal stages as the source of multiple cell types throughout the life of most animal species. In the mammalian forebrain, particularly in rodents, the ventricular-subventricular zone (V-SVZ) lining the lateral ventricles is a prominent postnatal stem cell niche capable of generating neuronal and glial cell progenies. The V-SVZ are the source of inhibitory neurons (interneurons) that migrate and integrate within the neural network of the olfactory bulb (OB) to influence behaviors including predator avoidance and sex-specific responses (Sakamoto *et al.*, 2011). The V-SVZ is composed of Type B cells, transit amplifying cells (TACs), Type A cells, and a monolayer of ependymal cells along the ventricular wall. Type B cells are the primary neural stem cells (NSC) of the V-SVZ, capable of generating TACs that divide into immature cell types that migrate into various forebrain structures where they mature (Lim and Alvarez-Buylla, 2016). Specifically, neurogenic TACs differentiate into immature neurons or Type A cells that migrate through the rostral migratory stream (RMS) and differentiate into molecularly distinct interneuron subtypes of the OB circuitry.

Studies of neurogenesis in the V-SVZ have expanded our knowledge on how diverse neuronal cell types are produced and integrate in the postnatal brain. However, the regulatory mechanisms that control stem/precursor behavior to establish the V-SVZ neurogenic niche at early neonatal stages are largely unclear. Control of stem/precursor behaviors at this early stage is critical to avoid early exhaustion of stem cell sources necessary for later adaptation and homeostatic needs, but also to avoid malformations and tumorigenesis. Therefore, further elucidation of the mechanisms controlling development of the V-SVZ, particularly the precise generation of specific stem/precursor cells at early neonatal stages, have far-reaching implications.

Suppressor of Fused (Sufu) is a cytoplasmic protein known for antagonizing Sonic hedgehog (Shh) signaling activity (Ramsbottom and Pownall, 2016). In the developing mammalian forebrain, Sufu modulates Shh signaling to control the proliferation, specification, and differentiation of radial glial progenitor cells (RGC) and their progenies (Yabut *et al.*, 2015, 2016; Yabut and Pleasure, 2018). Sufu exerts this role by controlling the activity of Gli transcription factors, the Shh signaling effectors, by proteolytic processing or protein stabilization to promote the repressor function of Gli and the eventual downregulation of Shh signaling target gene expression. Given the diverse roles of Sufu during corticogenesis in regulating RG cells, which eventually generate or transform to Type B cells in the postnatal brain, we wondered whether Sufu function in RG cells influence the generation of Type B cells and their progenies, and the establishment of the V-SVZ at neonatal stages. To investigate this, we employed a Cre-loxP approach to selectively delete Sufu in forebrain RGCs that give rise to Type B cells in the V-SVZ of the mouse forebrain. We found that loss of Sufu caused a dramatic cellular expansion in the dorsal V-SVZ by postnatal day (P) 7 of the neonatal mouse. This resulted in excess production of Sox2+ Type B cells and a subset of TACs. Specifically, loss of Sufu resulted in the expansion of ventral forebrain derived Gsx2+ TACs while simultaneously resulting in the dramatic decrease of dorsal forebrain derived Tbr2+ TACs. We found that these defects were partially due to a decrease in Gli3 expression, but not due to Shh signaling activation. Taken together, these studies indicate that Sufu plays a critical role in regulating neural cell precursor generation in the neonatal forebrain.

## EXPERIMENTAL PROCEDURES

### Animals

Mice carrying the floxed Sufu allele (Sufu^fl^) were kindly provided by Dr. Chi-Chung Hui (University of Toronto) and were genotyped as described elsewhere (Pospisilik *et al.*, 2010). The hGFAP-Cre (Stock #004600) was obtained from Jackson Laboratories (Bar Harbor, ME, USA). Mice designated as controls did not carry the *hGFAP*^*Cre*^ transgene and may have either one of the following genotypes: *Sufu*^*fl/+*^ or *Sufu*^*fl/fl*^ (**Figure 1A**). All animal protocols were in accordance to the National Institute of Health regulations and approved by the UCSF Institutional Animal Care and Use Committee (IACUC).

**Figure 1.**
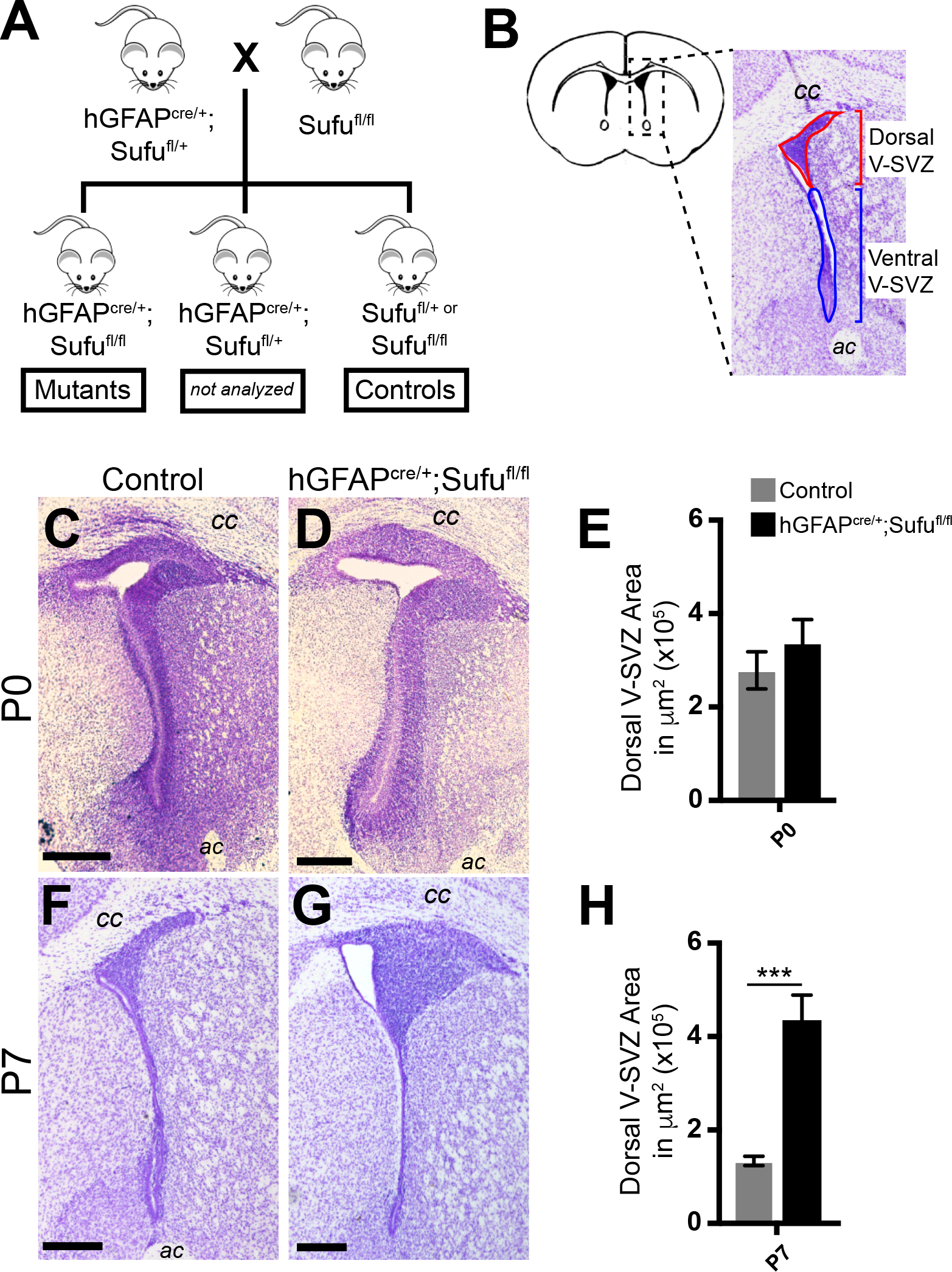
Loss of SUFU causes an expansion of dorsal V-SVZ cells at early postnatal stages. **(A)** Illustration of the breeding scheme used to generate conditional Sufu knockouts and controls for analysis. **(B)** Schematic diagram of the dorsal and ventral V-SVZ areas analyzed in these studies. **(C, D)** Cresyl-violet staining of coronal sections of the P0 *hGFAP*^*cre/+*^*;Sufu*^*fl/fl*^ and control littermates show no anatomical or structural difference in V-SVZ between the two genotypes. **(E)** Quantification of V-SVZ area showing no significant difference between the size of the V-SVZ between P0 *hGFAP*^*cre/+*^*;Sufu*^*fl/fl*^ mice and controls. **(F, G)** Cresyl-violet staining show an obvious increase in size of the dorsal V-SVZ of the P7 *hGFAP*^*cre/+*^*;Sufu*^*fl/fl*^ mice compared to controls. **(H)** Quantification of V-SVZ area shows a nearly three-fold increase in the mutant dorsal V-SVZ compared to controls. *** p-value ≤ 0.01; V-SVZ, ventricular-subventricular zone. P0, postnatal day 0; P7, postnatal day 7. ac, anterior commissure; cc, corpus callosum

### Quantitative PCR

Total RNA was isolated from dissected V-SVZ of P7 mice using TRIzol™ Reagent (Thermo Fishire Scientific), according to the manufacturer’s instructions, and each sample was reverse-transcribed using a SuperScript IV cDNA Synthesis Kit (Invitrogen). Quantitative PCR reactions were performed using a KAPA SYBR Fast qPCR Kit (KAPA Biosystems) with ROX as reference dye, and transcript expression was measured via Applied Biosystem 7500 Real-Time PCR System (Life Technologies). Expression levels of each gene were normalized to RNA polymerase II subunit A (polr2a) and calculated relative to the control. The following primers were used: Gli1 Fw: CCGACGGAGGTCTCTTTGTC; Gli1 Rv AACATGGCGTCTCAGGGAAG; Gli3 Fw: AAGCGGTCCAAGATCAAGC; Gli3 Rv: TTGTTCCTTCCGGCTGTTC; Ptch1 Fw:TGACAAAGCCGACTACATGC; Ptch1 Rv:AGCGTACTCGATGGGCTCT; Polr2a Fw: CATCAAGAGAGTGCAGTTCG; Polr2a Rv: CCATTAGTCCCCCAAGTTTG.

### Immunohistochemistry and BrdU-Labeling

Perfusion, dissection, immunofluorescence, and Nissl staining were conducted according to standard protocols as previously described (Siegenthaler *et al.*, 2009). Cryostat sections were air dried and rinsed 3× in PBS plus 0.2%Triton before blocking for 1 h in 10% normal lamb serum diluted in PBS with 0.2% Triton to prevent nonspecific binding. Primary antibodies were diluted in 10% serum diluted in PBS with 0.2% Triton containing 40,6-diamidino-2-phenylindole (DAPI); sections were incubated in primary antibody overnight at room temperature. The following antibodies were used: mouse anti-BrdU (1:50 dilution; BD Pharmingen, Franklin Lakes, NJ, USA), rabbit anti-Phospho-Histone H3 (1:250 dilution; Millipore, Billerica, MA, USA), rabbit anti-Sox2 (1:1000 dilution; Abcam, Cambridge, UK), rabbit anti-Tbr2 (1:500 dilution; Abcam, Cambridge, UK), mouse anti-Tuj1 (1:500 dilution; Covance, Princeton, NJ, USA), rabbit anti-Cleaved Caspase 3 (1:300 dilution; Cell Signaling, Madison, WI, USA), mouse anti-Calretinin (1:250 dilution; Millipore, Billerica, MA, USA), mouse anti-Ki67(1:2500 dilution; Millipore, Billerica, MA, USA), rabbit anti-GSX2 (1:250 dilution; gift from Kenneth Campbell (Toresson, Potter and Campbell, 2000)) mouse anti-Olig2 (1:250 dilution; Millipore, Billerica, MA, USA), rabbit anti-Calbindin (1:1000 dilution; Swant), rabbit anti-Pdgfra (1:1000; gift from William Stallcup (Nishiyama *et al.*, 1996)) rabbit anti-Doublecortin (1:250 dilution; Abcam, Cambridge, UK);. For 5-bromo-2-deoxyuridine (BrdU, Sigma, St. Louis, MO, USA) labeling, early postnatal mice were treated with 50 μg/g BrdU by intraperitoneal injection at P0-P1 prior to dissection at P7-P8. For BrdU-labelling at P28, mice were treated with 50 μg/g BrdU pulse by intraperitoneal injection from P6 to P7 every 12 hours for 36 hours for a total of 3 BrdU treatments. To detect primary antibodies, we used species-specific Alexa Fluor-conjugated secondary antibodies (1:500; Invitrogen) in 1X PBS-T for 1 h at room temperature, washed with 1X PBS, and coverslipped with Fluoromount-G (SouthernBiotech).

### Image Analysis and Acquisition

Images were acquired using a Nikon E600 microscope equipped with a QCapture Pro camera (QImaging) or Zeiss Axioscan Z.1 (Zeiss, Thornwood, NY, USA) using the Zen 2 blue edition software (Zeiss, Thornwood, NY, USA). NIH Image J was used to quantify raw, unedited images. All analyses were conducted in at least 2-3 20 μm thick sections that were histologically matched at the rostral-caudal level between genotypes.

#### V-SVZ Analysis

For measurement of VZ/SVZ thickness, the length of densely populated cell or DAPI+ regions adjacent to the lateral ventricles was measured and designated as the dorsal V-SVZ or ventral V-SVZ as defined in **Figure 1B**. DAPI-dense regions were also used to define and measure the SVZ, RMS, and the OB to quantify BrdU-localization along sagittal sections. Cells labeled with cell-specific markers were quantified within the dorsal V-SVZ to measure the number of cells per 100 μm^2^ or per V-SVZ.

#### Olfactory Bulb (OB) Analysis

A slice of the OB containing all layers, as designated according to their anatomical features (as defined in **Figure 7B**), was used for cell quantification. Cells that express cell-specific markers (Calretinin+, Calbindin+, or BrdU+) were counted in each layer.

### Statistics

All experiments were conducted in triplicate with a sample size of n = 3-6 embryos/animals per genotype. Unpaired Student t-test was conducted using Prism 7 (GraphPad) for pairwise analysis of control and mutant genotypes. Graphs display the mean ± standard error of the mean (SEM).

## RESULTS

### Expansion of the V-SVZ in the P7 *hGFAP*^*cre/+*^*;Sufu*^*fl/fl*^ mice

Type B cells of the V-SVZ are produced at late embryonic stages from RGCs of the embryonic dorsal and ventral forebrain. To ensure deletion of Sufu Type B cells, we utilized the hGFAP-Cre mouse line in which Cre Recombinase is specifically active in RG cells of the late stage embryonic forebrain, at E13.5 (Zhuo *et al.*, 2001). Generating the *hGFAP*^*cre/+*^*;Sufu*^*fl/fl*^ mice (**Figure 1A**) allowed us to target Sufu deletion in RGCs from all progenitor domains of the dorsal and ventral forebrains. At P0, we examined coronal sections from V-SVZ regions of *hGFAP*^*cre/+*^*;Sufu*^*fl/fl*^ and control littermates and found no obvious anatomical and structural differences in the V-SVZ between these two genotypes (**Figure 1C-1D**). By P7, we found a dramatic enlargement of the dorsal V-SVZ in mutant mice compared to control littermates (**Figure 1F-1G**). Quantification of the overall dorsal V-SVZ area confirmed that no significant difference in the overall size of the dorsal V-SVZ (**Figure 1B**) was observed between controls and mutants at P0 (**Figure 1E**; 278,512 ± 39,546 μm^2^ for n = 3 controls and 338,946 ± 48,133 μm^2^ for n = 3 mutants; *p*-value = 0.369). However, the mutant dorsal V-SVZ was expanded approximately three-fold compared to control littermates by P7 (**Figure 1H**; 126,984 ± 9,915 μm^2^ for n=4 controls and 514,863 ± 86,674 μm^2^ for n=4 mutants; p-value =0.0043). These observations indicate that loss of Sufu results in uncontrolled cell expansion specifically in the dorsal V-SVZ at early neonatal stages.

### Accumulation of proliferating cells in the dorsal V-SVZ of the P7 *hGFAP*^*cre/+*^*;Sufu*^*fl/fl*^ mice

The dorsal V-SVZ is populated by actively proliferating precursors, including immature Type A cells that divide and migrate into the OB. To examine whether the increase in cell number in the P7 *hGFAP*^*cre/+*^*;Sufu*^*fl/fl*^ dorsal V-SVZ is due to the failed migration of Type A cells, we labeled proliferating precursors in the V-SVZ of P0/P1 littermates by intraperitoneal injection of 5-bromo-2-deoxyuridine (BrdU) and examined the location of BrdU-labeled (BrdU+) cells at P7 (**Figure 2A**). Proliferating cells in the dorsal V-SVZ Cells will be labeled with BrdU at P1 and include TACs destined to differentiate into Type A cells that will subsequently migrate anteriorly through the RMS and finally to the OB. Thus, we were able to trace the location of BrdU+ cells along this migratory route and determine whether labeled migratory Type A cells successfully reached the OB. The location of BrdU+ cells was examined in sagittal sections of P7 brains. As expected, BrdU+ cells were observed in the V-SVZ, the RMS, and the OB of control mice, indicating that BrdU-labeled V-SVZ cells at P1 successfully migrated into the OB by P7 (**Figure 2B-2C**). Similarly, we found BrdU+ cells in the V-SVZ, RMS, and OB of P7 *hGFAP*^*cre/+*^*;Sufu*^*fl/fl*^ brains (**Figure 2D-2E**). However, an obvious increase in BrdU+ cells were observed in the P7 *hGFAP*^*cre/+*^*;Sufu*^*fl/fl*^ dorsal V-SVZ (arrow, **Figure 2E**) but not in controls (arrow, **Figure 2C**). **Figure 2F** shows that quantification of BrdU+ cells resulted in a significant increase in the P7 *hGFAP*^*cre/+*^*;Sufu*^*fl/fl*^ V-SVZ compared to controls (**Figure 2F**; 0.1154 ± 0.01794 cells per 100 μm^2^ for n=3 controls and 0.2183 ± 0.02015 cells per 100 μm^2^ for n=3 mutants; p-value=0.0189), whereas no significant difference were quantified in the RMS (**Figure 2F**; 0.1308 ± 0.01477 cells per 100 μm^2^ for n=3 controls and 0.1789 ± 0.03221 cells per 100 μm^2^ for n=3 mutants; p-value=0.2463) and OB (**Figure 2F**; 0.1225 ± 0.002195 cells per 100 μm^2^ for n=3 controls and 0.1457 ± 0.01775 cells per 100 μm^2^ for n=3 mutants; p-value=0.2650) between controls and mutants. Overall, these observations indicate that cells within the dorsal V-SVZ of mutant mice were able to migrate despite of the accumulation of BrdU+ cells in the mutant dorsal V-SVZ.

**Figure 2:**
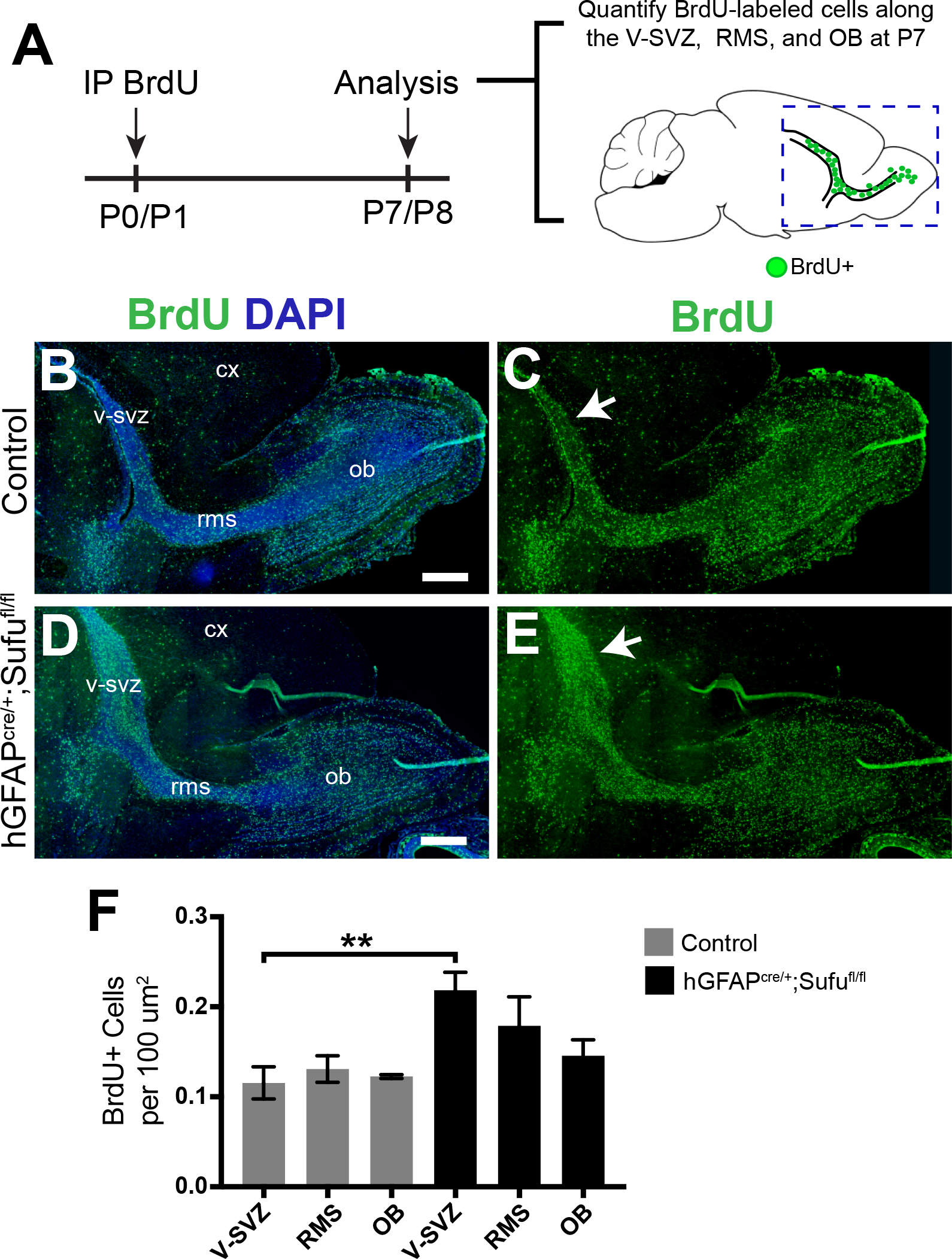
Cells generated at neonatal stages accumulate in the dorsal V-SVZ of the P7 *hGFAP*^*cre/+*^*;Sufu*^*fl/fl*^ mice. **(A)** Schematic of BrdU-labeling experiments. Intraperitoneal injections of S-label, 5-bromo-2-deoxyuridine (BrdU), were administered to P0/P1 littermates and quantification of actively proliferating cells in the V-SVZ, RMS, and OB of sagittal sections were performed at P7. (B-E) Immunofluorescence staining with anti-BrdU shows successful migration of actively proliferating progenitors from the V-SVZ through the RMS, and into the OB of P7 *hGFAP*^*cre/+*^*;Sufu*^*fl/fl*^ mice and control littermates. Arrows indicate an observable increase in BrdU+ cells in the P7 *hGFAP*^*cre/+*^*;Sufu*^*fl/fl*^ dorsal V-SVZ compared to controls. (F) Quantification confirms a significant increase in BrdU+ cells in the P7 *hGFAP*^*cre/+*^*;Sufu*^*fl/fl*^ dorsal V-SVZ compared to controls whereas no significant differences in the number of BrdU+ cells were observed in the RMS or OB. ** p-value ≤ 0.01; V-SVZ, ventricular-subventricular zone; RMS, rostral migratory stream; OB, olfactory bulb; CX, cortex

### Persistent cell proliferation in the dorsal V-SVZ of the P7 *hGFAP*^*cre/+*^*;Sufu*^*fl/fl*^ mice

NSCs in the V-SVZ include slowly-dividing quiescent populations that are able to retain S-phase labels such as BrdU, referred to as label-retaining cells (LRC), for extended periods (Cotsarelis, Sun and Lavker, 1990; Codega *et al.*, 2014). To exclude the possibility that BrdU+ cells in the dorsal V-SVZ are label-retaining quiescent NSCs (qNSC), we examined the proportion of cells that remained proliferative by immunostaining with the mitotic marker Phospho-Histone H3 (PH3) after prolonged survival following BrdU labeling (**Figure 3A**). We found that unlike controls (**Figure 3B**), many of the accumulated BrdU+ cells in the P7 *hGFAP*^*cre/+*^*;Sufu*^*fl/fl*^ dorsal V-SVZ expressed PH3, and were therefore still proliferative (**Figure 3C**). Quantification of double-labeled cells verified that a significantly higher proportion of PH3+ and BrdU+ double-labeled cells were present in the P7 *hGFAP*^*cre/+*^*;Sufu*^*fl/fl*^ dorsal V-SVZ (**Figure 3D**, 0.003417 ± 0.0004118 cells per 100 μm^2^ for n=3 controls and 0.007374 ± 0.001042 cells per 100 μm^2^ for n=3 mutants; p-value= 0.0242). Our findings indicated that loss of Sufu resulted in the continuous proliferation of cells within the dorsal V-SVZ of the P7 *hGFAP*^*cre/+*^*;Sufu*^*fl/fl*^ mice.

**Figure 3:**
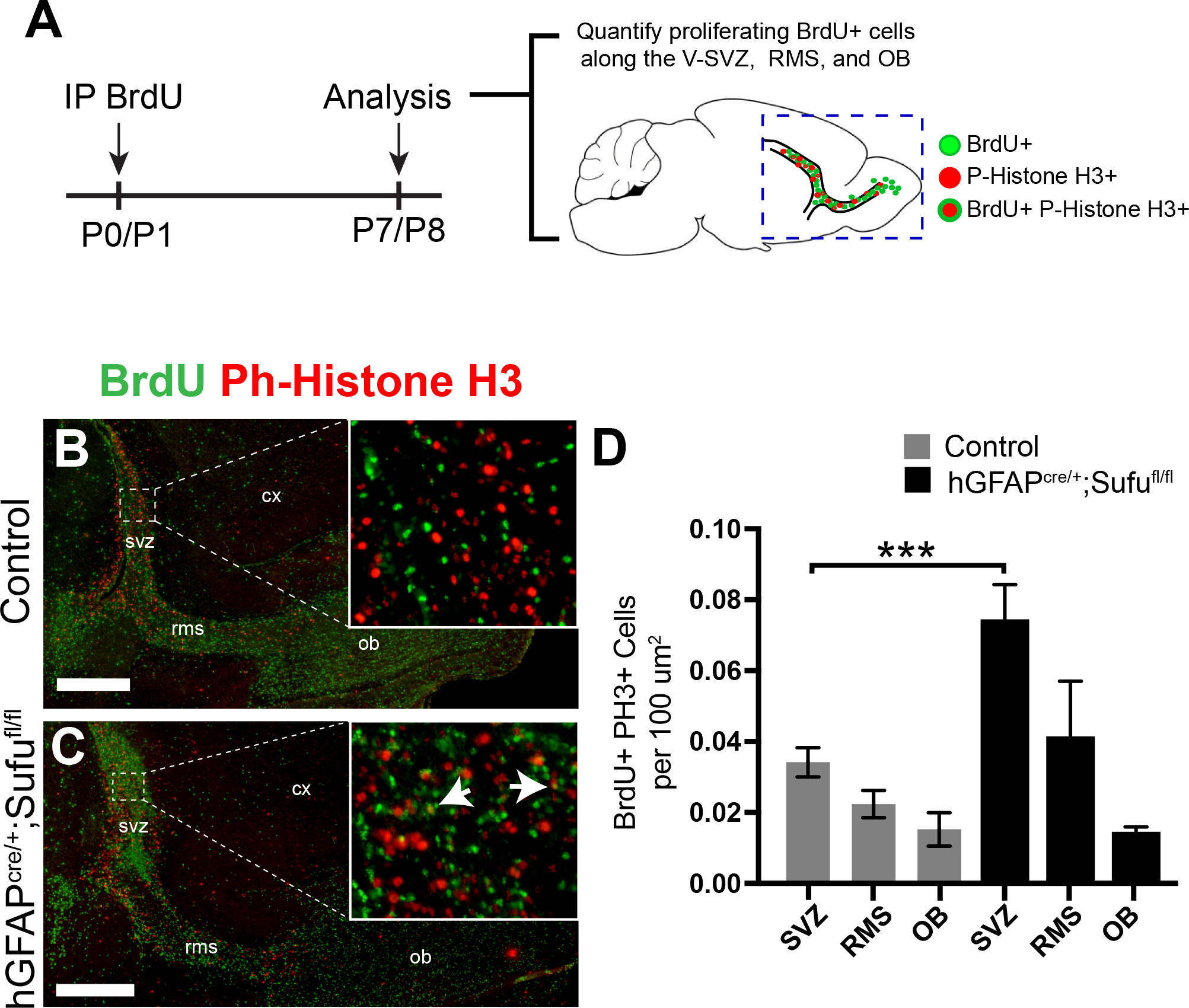
Persistent cell proliferation in the dorsal V-SVZ of the P7 *hGFAP*^*cre/+*^*;Sufu*^*fl/fl*^ mice. **(A)** Schematic of BrdU-labeling experiments to identify proliferating cells. Intraperitoneal injections of S-phase label, 5-bromo-2-deoxyuridine (BrdU), were administered to P0/P1 littermates and quantification of double-labeled BrdU+ and P-Histone H3+ cells in the V-SVZ, RMS, and OB of sagittal sections were performed at P7. **(B,C)** Immunofluorescence staining with anti-P-Histone H3, a mitotic marker, and anti-BrdU shows a visible increase in BrdU+ and Phospho-histone H3+ double-labeled cells (arrowheads) in the dorsal V-SVZ of P7 *hGFAP*^*cre/+*^*;Sufu*^*fl/fl*^ mice compared to controls. **(D)** Quantification of BrdU+ and P-Histone H3+ double-labeled cells verified an increase in the dorsal V-SVZ of the P7 *hGFAP*^*cre/+*^*;Sufu*^*fl/fl*^ mice compared to controls, however, no significant increase was quantified in the RMS or OB of P7 *hGFAP*^*cre/+*^*;Sufu*^*fl/fl*^ mice compared to controls. ***p-value ≤ 0.01; V-SVZ, ventricular-subventricular zone; RMS, rostral migratory stream; OB, olfactory bulb

### Loss of Sufu drives the proliferation of Type B cells in the dorsal V-SVZ

In the dorsal V-SVZ, the transcription factors Gli2 and Gli3, typically under Sufu control, are specifically expressed in Type B cells of the dorsal V-SVZ (Petrova, Garcia and Joyner, 2013; Lin *et al.*, 2014). To examine whether the increase in proliferation specifically affected B cells, we conducted immunostaining with Sox2, a transcription factor highly expressed in V-SVZ Type B cells (Ellis *et al.*, 2004). Indeed, we observed an increase in cells highly expressing Sox2 (Sox2+) in the P7 *hGFAP*^*cre/+*^*;Sufu*^*fl/fl*^ dorsal V-SVZ, particularly in regions where Sox2+ cells were typically scant in the controls (boxed areas, **Figure 4A-4D**). Many of these co-expressed Nestin demonstrating that these Sox2+ cells are Type B cells (arrowheads, Figure 4D). When quantified, we did not find any significant difference per unit area in the number of Sox2+ cells (**Figure 4E**, 0.6895 ± 0.04573 cells per 100 μm^2^ for n=3 controls and 0.5037 ± 0.0773 cells per 100 μm^2^ for n=3 mutants; p-value=0.1074). However, we found that the total number of Sox2+ cells dramatically increased per dorsal V-SVZ of mutant mice (**Figure 4F**, 351.8 ± 59.82 cells, n=3 controls and 1521 ± 391.5 cells, n=3 mutants; p-value=0.0418). These findings showed that Sox2+ Type B cells proliferated proportionately as the dorsal V-SVZ expanded. To examine whether Sox2+ NSCs were proliferative, we scored for Sox2+ and Ki67+ cells. Unlike in controls, there was a significant increase in double-labeled Sox2+ and Ki67+ cells in the dorsal V-SVZ of mutant mice (**Figure 4G-4I**; 24.13 ± 3.21 cells, n=4 controls; 178.6 ± 59.11 cells, n=4 mutants; p-value= 0.0401). Furthermore, when compared to controls, a significant percentage of Sox2+ cells were Ki67+ (**Figure 4J**; 7.472 ± 1.141% cells n=4 controls; 13.12 ± 0.9569%, n=4 mutants; p-value = 0.0091). Thus, loss of Sufu in Type B cells promoted their uncontrolled proliferation contributing significantly to the expansion of the dorsal V-SVZ.

**Figure 4:**
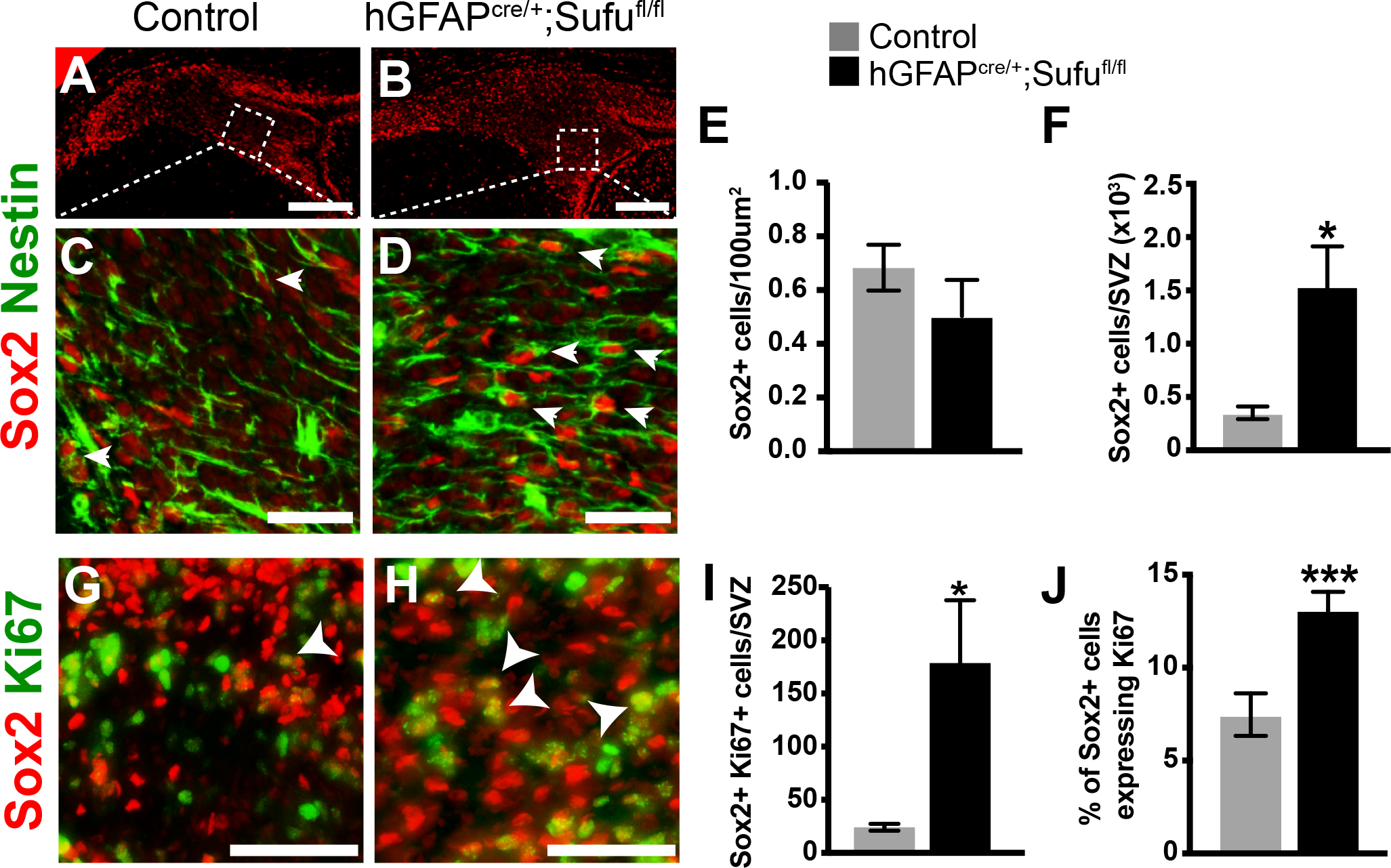
Loss of Sufu drives the proliferation of Type B cells in the dorsal V-SVZ. **(A, D)** Immunofluorescence staining with Type B cell marker, Sox2, shows an increase in Sox2+ cells in the dorsal V-SVZ of P7 *hGFAP*^*cre/+*^*;Sufu*^*fl/fl*^ mice (B) compared to controls (A). Boxed inset pertains to high magnification images in C and D, Scale bars = 250 m. (C, D) Double-immunofluorescence staining with Sox2 and Nestin show that cells with high levels of Sox2 expression in the control (C) and mutant dorsal V-SVZ (D) also co-express the neural stem cell marker, Nestin, and are therefore multipotent Type B cells. Notably, many of the Sox2+/Nestin+ double-labeled cells in the mutant dorsal V-SVZ is found beyond the ventricular wall. Scale bars = 50 5m. Arrowheads mark double-labeled cells. **(E, F)** Quantification of Sox2+ cells in the dorsal V-SVZ shows no significant differences per 100 um^2^ (E) but showed a significant increase in the number of Sox2+ cells per dorsal V-SVZ (F) in the P7 *hGFAP*^*cre/+*^*;Sufu*^*fl/fl*^ mice, indicating that Sox2+ cells increased in proportion to the expanding dorsal V-SVZ of mutant mice. **(G, H)** Immunofluorescence staining against the proliferation marker, Ki67, and Type B cell marker, Sox2, show a visible increase in the number of double-labeled cells of the P7 *hGFAP*^*cre/+*^*;Sufu*^*fl/fl*^ dorsal V-SVZ (G) compared to controls (H). Scale bars = 50 5m. Arrowheads mark double-labeled cells. **(I, J)** Quantification of double-labeled Ki67+ and Sox2+ cells verify a significant increase in the density (I) and percentage (J) of proliferating Sox2+ cells in the P7 *hGFAP*^*cre/+*^*;Sufu*^*fl/fl*^ dorsal V-SVZ compared to controls. * p-value ≤ 0.05; ***p-value ≤ 0.01

### Differential effects on transit amplifying cells in the dorsal V-SVZ of P7 *hGFAP*^*cre/+*^*;Sufu*^*fl/fl*^ mice

Since Type B cells in the dorsal V-SVZ generate neurogenic TACs, we wondered whether the increase in proliferating Sox2+ cells indicated that these cells were maintained in their multipotent state, failing to generate more mature precursor cell types, such as TACs. We therefore sought to determine the number of TACs by immunostaining with either Gsx2 or Tbr2 expression. Our results showed that both Gsx2+ and Tbr2+ TACs were present in the dorsal V-SVZ of P7 *hGFAP*^*cre/+*^*;Sufu*^*fl/fl*^ mice, but drastically differed in numbers. Visibly, Gsx2+ cells appeared less in the mutant dorsal V-SVZ compared to controls (**Figure 5A-5B**). However, quantification of Gsx2+ cells revealed no significant difference in the density of Gsx2+ cells between control and mutant dorsal V-SVZ (**Figure 5C**, 0.02794 ± 0.007368 cells per 100 μm^2^ for n=3 controls and 0.02412 ± 0.006588 cells per 100 μm^2^ for n=3 mutants; p-value=0.7188). We also found a significant increase in the number of Gsx2+ cells per dorsal V-SVZ in the mutant mice (**Figure 5D**, 85.17 ± 15.58 cells per SVZ for n=3 controls; 155 ± 4.368 cells per SVZ for n=3 mutants; p-value = 0.0125), indicating that the number of Gsx2+ cells increased in proportion to the expanding dorsal V-SVZ. Further analysis showed that the proliferative capacity of Gsx2+ TACs did not significantly change since the proportion of Gsx2+ TACs expressing Ki67 were comparable between mutant and control dorsal V-SVZ (**Figure 5G**, 23.24 ± 6.655% of cells in controls, 28.62 ± 4.27% of cells in mutants; n=3 control/mutant mice; p-value 0.5337). In contrast, Tbr2+ TACs were markedly reduced in the dorsal V-SVZ of mutant mice (**Figure 5H-5I**). Quantification of Tbr2+ cells showed a significant reduction in the density (0.1629 ± 0.01479 cells per 100 μm^2^ for n=3 controls and 0.0128 ± 0.003706 cells per 100 μm^2^ for n=3 mutants; p-value=0.0006) and total number (76.67 ± 5.069 cells/SVZ for n=3 controls and 47.78 ± 6.859 cells/SVZ for n=3 mutants; p-value=0.0276) in the dorsal V-SVZ of P7 *hGFAP*^*cre/+*^*;Sufu*^*fl/fl*^ mice compared to controls (**Figure 5J-5K**). To examine whether the reduction in Tbr2+ cells was due to the inability to proliferate, we double-labeled Tbr2+ cells with the proliferation marker Ki67. As with controls (**Figure 5L**), Tbr2+ cells in the mutant dorsal V-SVZ co-labeled with Ki67 (**Figure 5M**), indicating that the Tbr2+ cells were able to proliferate. Our quantification verified these observations in which no significant difference in the number of proliferating Tbr2+ cells between mutants and controls was evident (**Figure 5N**; 42.8 ± 3.811% of Tbr2+ cells in controls and 35.95 ± 3.483% of Tbr2+ cells in mutants, p-value = 0.2554; n=3/genotype).

**Figure 5.**
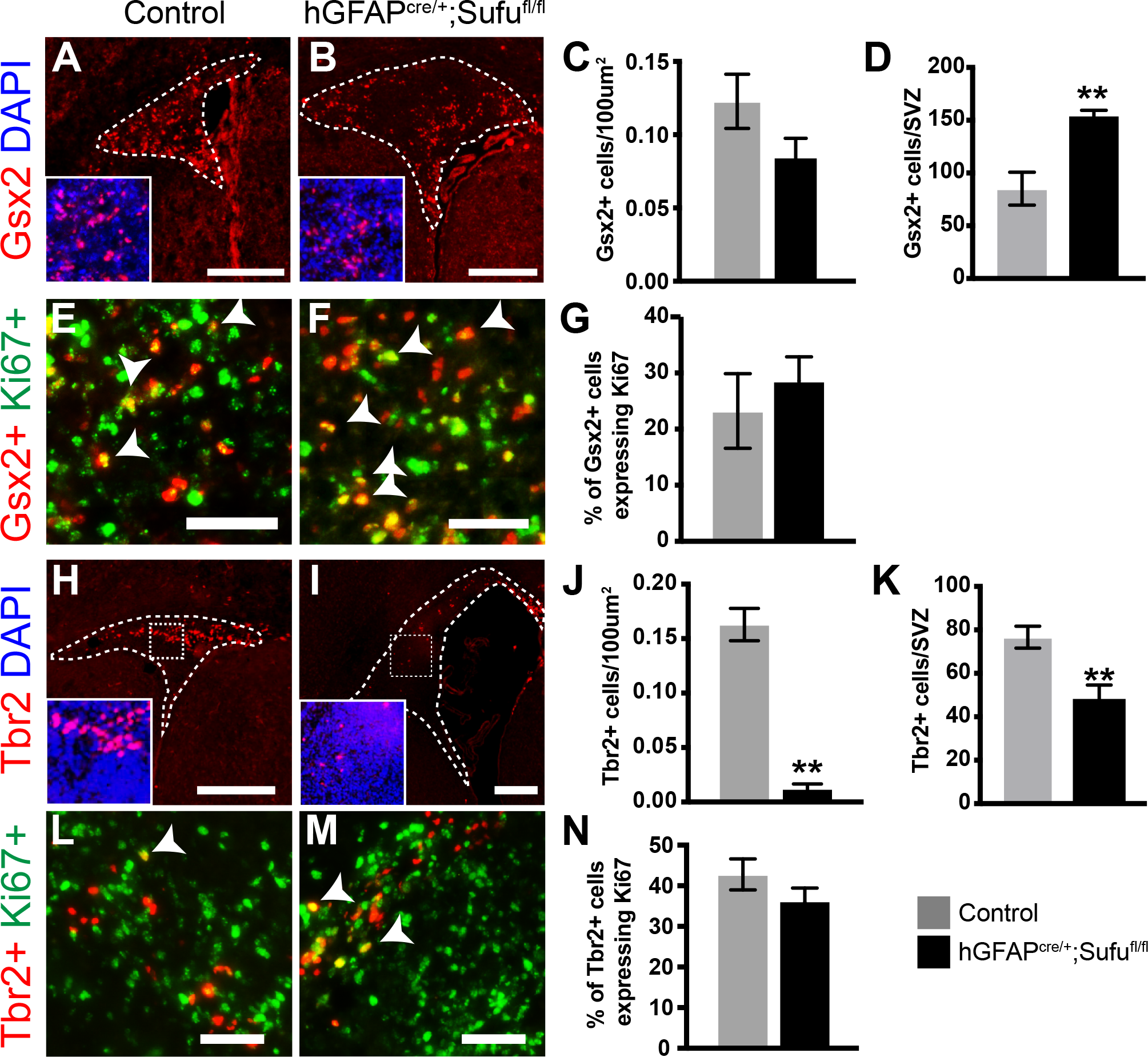
Differential effects of Sufu deletion in the proliferation of transit amplifying cells (TACs) in the P7 *hGFAP*^*cre/+*^*;Sufu*^*fl/fl*^ dorsal V-SVZ. **(A, B)** Immunofluorescence staining against Gsx2, a marker for TACs, show that Gsx2+ cells are present in both control **(A)** and mutant **(B)** P7 dorsal V-SVZ where Gsx2+ cells appear to be reduced in density. Scale bars = 250 2m **(C, D)** Quantification of Gsx2+ cells in the P7 dorsal V-SVZ shows a slight, but insignificant, reduction in the density of Gsx2+ cells in the mutant mice (C). On the other hand, the number of Gsx2+ cells in the dorsal V-SVZ of mutant mice are significantly increased compared to controls (D). **(E, F)** Double immunofluorescence staining against Ki67 and Gsx2, show a visible increase in the number of double-labeled cells of the P7 *hGFAP*^*cre/+*^*;Sufu*^*fl/fl*^ dorsal V-SVZ. Scale bars = 50 5m. **(G)** Quantification of proliferating Gsx2+ cells, co-labeled with the proliferation marker Ki67, verify that Gsx2+ cells in the P7 *hGFAP*^*cre/+*^*;Sufu*^*fl/fl*^ dorsal V-SVZ did not proliferate at higher rates compared to controls. **(H, I)** Immunofluorescence staining against dorsal forebrain neurogenic progenitor cell marker, Tbr2, shows a visible decrease in the number of Tbr2+ cells in the P7 *hGFAP*^*cre/+*^*;Sufu*^*fl/fl*^ dorsal V-SVZ (F) compared to controls (E). Scale bars = 250 μm. **(J, K)** Quantification of Tbr2+ cells in the dorsal V-SVZ shows a significant decrease in the density (J) and overall number (K) in the dorsal V-SVZ in P7 *hGFAP*^*cre/+*^*;Sufu*^*fl/fl*^ mice indicating an overall decrease in Tbr2+ precursors. **(L, M)** Double immunofluorescence staining against Ki67 and the precursor cell marker, Tbr2, show proliferating Tbr2+ cells in both control (L) and mutant (M) P7 dorsal V-SVZ. Scale bars = 50 5m. Arrowheads mark double-labeled cells. **p-value ≤ 0.03. **(N)** Quantification of proliferating Tbr2+ cells, co-labeled with the proliferation marker Ki67, show no significant differences in the number of proliferating cells between control and mutant P7 dorsal V-SVZ.

Taken together, our observations showed that Tbr2+ and Gsx2+ TACs did not proliferate at higher rates in the dorsal V-SVZ of P7 *hGFAP*^*cre/+*^*;Sufu*^*fl/fl*^ mice compared to controls. This suggested that the reduction of Tbr2+ TACs in the mutant dorsal V-SVZ was caused by the failure of Type B cells to generate these precursors. Rather, Type B cells in the dorsal V-SVZ of mutant mice preferentially generated Gsx2+ TACs. Given that Gsx2+ and Tbr2+ are derived from NSCs that originate from ventral and dorsal forebrain regions, respectively (Kowalczyk *et al.*, 2009; López-Juárez *et al.*, 2013), these observations reveal that Sufu differentially affects subpopulations of Type B cells that are molecularly distinct according to their embryonic origins.

### Loss of Suppressor of Fused results in downregulated V-SVZ Gli3 expression independent of Shh signaling

Similar to the P7 *hGFAP*^*cre/+*^*;Sufu*^*fl/fl*^ V-SVZ, deletion of Gli3 in NSCs of the developing brain cause expansion of the dorsal V-SVZ in neonatal mice (Petrova, Garcia and Joyner, 2013; Wang *et al.*, 2014). We examined whether changes in Gli3 levels occurred in the P7 *hGFAP*^*cre/+*^*;Sufu*^*fl/fl*^ V-SVZ. Quantitative PCR analysis showed that Gli3 mRNA was reduced in dissected V-SVZ of P7 *hGFAP*^*cre/+*^*;Sufu*^*fl/fl*^ mice (**Figure 6A**; 1 ± 0.09995 relative expression level in controls, 0.4697 ± 0.02742 levels in mutants, n=3 mice/genotype, p-value = 0.0069). These findings suggest that loss of Sufu results in diminished expression of Gli3 causing the defects observed in the *hGFAP*^*cre/+*^*;Sufu*^*fl/fl*^ mice. As with mice lacking Gli3 in the V-SVZ (Wang *et al.*, 2014), we did not observe ectopic activation of Shh signaling in the dorsal V-SVZ of the P7 *hGFAP*^*cre/+*^*;Sufu*^*fl/fl*^ mice by visualizing cells that express LacZ under the control of the Gli1 promoter (LacZ+) (Ahn and Joyner, 2005). Fewer LacZ+ cells were observed in the dorsal V-SVZ of mutant mice (**Figure 6C**), showing that proliferation of Shh-responsive cells did not increase in proportion to other cell types. This observation is likely the result of reduced Gli1 expression in the dorsal V-SVZ of mutant mice (**Figure 6B**). Additionally, these results showed that the expansion of the dorsal V-SVZ is not due to activation of Shh signaling in the neonatal P7 *hGFAP*^*cre/+*^*;Sufu*^*fl/fl*^ mice. Further supporting these observations, we also found by qPCR analysis that expression of the Shh target gene, Ptch1, was also comparable between control and mutant mice (**Figure 6B**).

**Figure 6:**
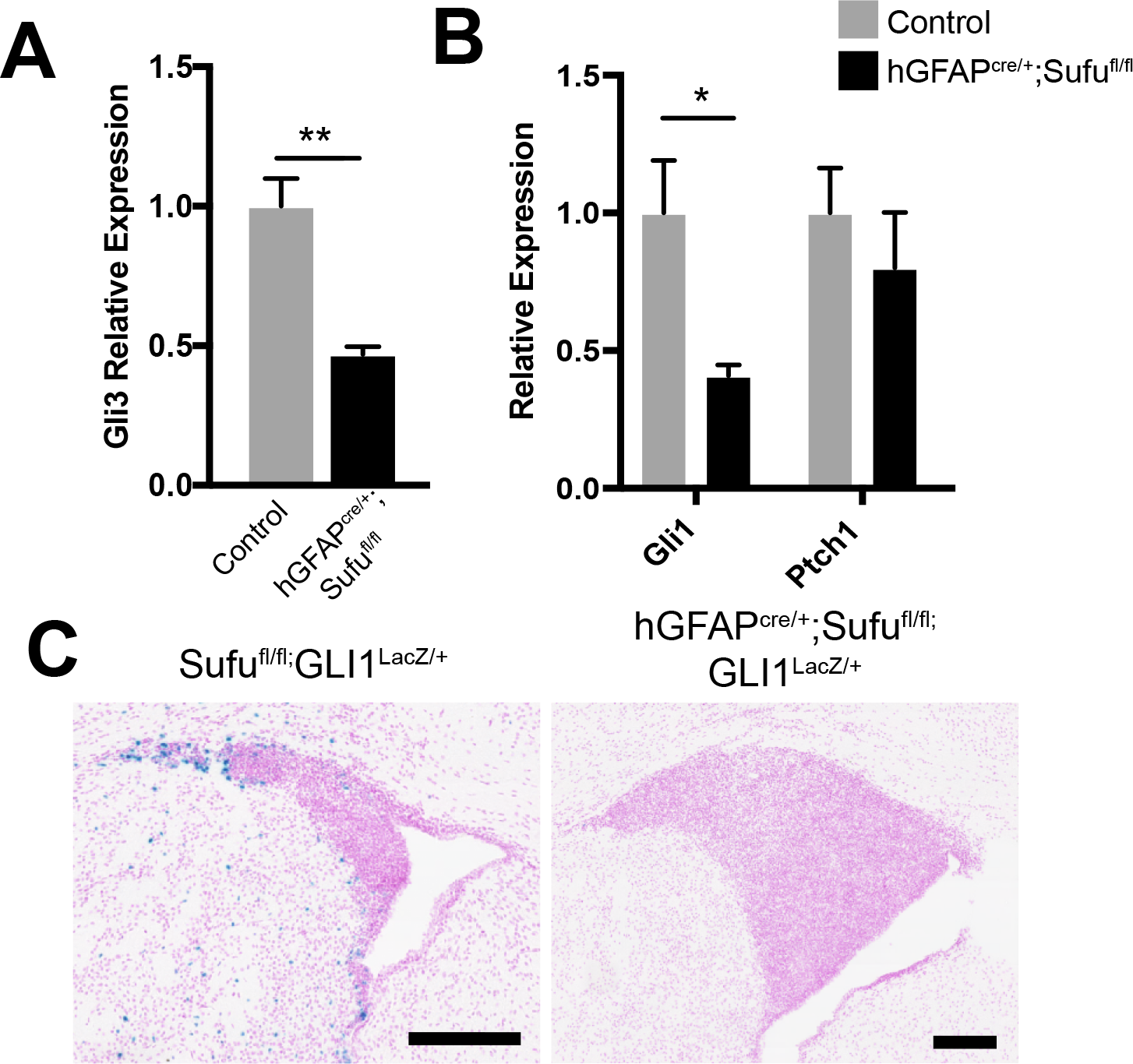
Loss of Suppressor of Fused results in downregulated Gli3 expression in the dorsal V-SVZ and did not ectopically activate Shh signaling. **(A)** qPCR analysis of Gli3 mRNA levels extracted from dissected V-SVZ show that Gli3 expression is significantly reduced in the P7 V-SVZ of *hGFAP*^*cre/+*^*;Sufu*^*fl/fl*^ mice. **(B)** qPCR analysis of Shh targets, Gli1 and Ptch1, show that Gli1 expression is significantly reduced in the P7 V-SVZ of *hGFAP*^*cre/+*^*;Sufu*^*fl/fl*^ mice compared to controls. However, levels of Ptch1 expression were comparable between controls and mutant mice, indicating that Shh signaling activity is not increased in the dorsal V-SVZ of mice lacking Sufu. **(C)** Shh-responsive cells, as detected by β-galactosidase activity (LacZ+) in mice carrying the Shh-reporter Gli1-LacZ, are largely absent in the dorsal V-SVZ of P7 *hGFAP*^*cre/+*^*;Sufu*^*fl/fl*^ mice whereas few LacZ+ cells were observed in the controls. Scale bars = 200 2m. * p-value ≤ 0.05; **p-value ≤ 0.03

**Figure 7.**
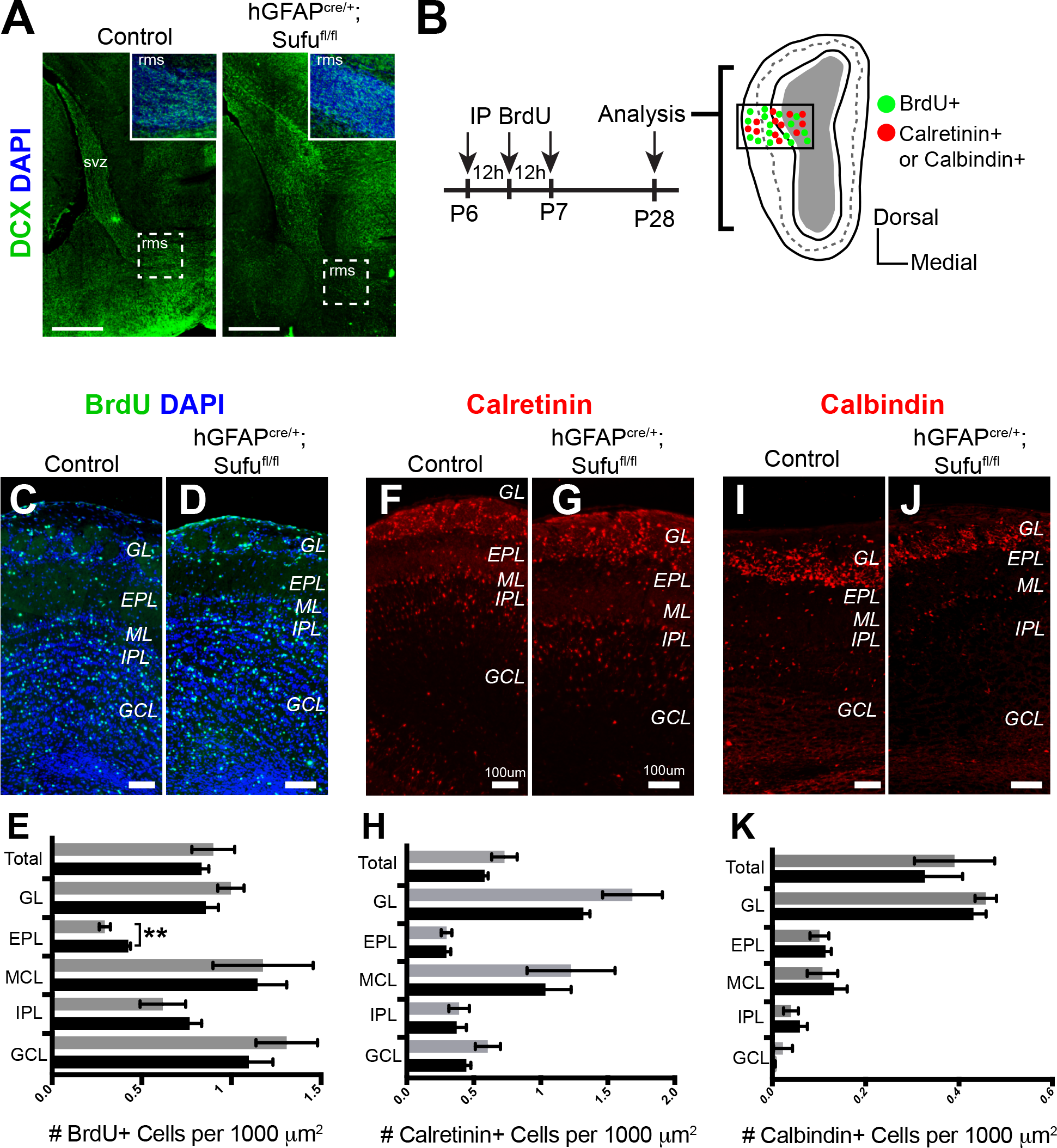
Type A cells are produced and able to differentiate into interneuron subtypes in the P28 *hGFAP*^*cre/+*^*;Sufu*^*fl/fl*^ olfactory bulb (OB). **(A)** Immunofluorescence staining against DCX, which labels Type A cells, indicates that Type A cells are generated by Gsx2+ or Tbr2+ neurogenic precursors and are able to migrate through the RMS. RMS, rostral migratory stream; OB, olfactory bulb; CX, cortex **(B)** Schematic of the experimental design to identify the progeny of proliferating cells in the P7 dorsal V-SVZ. Three pulsed intraperitoneal injections at 8-hour intervals of S-phase label, 5-bromo-2-deoxyuridine (BrdU), were administered starting at P6 to label proliferating cells in the V-SVZ and determine their localization and identity in the OB at P28. **(A)** Immunofluorescence staining against DCX, which labels Type A cells, indicates that Type A cells are generated by Gsx2+ or Tbr2+ neurogenic precursors and are able to migrate through the RMS. **(C, D)** Immunofluorescence staining against BrdU shows visible confirmation of proliferating type A cells migrating into the OB layers of P28 *hGFAP*^*cre/+*^*;Sufu*^*fl/fl*^ mice and controls. **(E)** Quantification of BrdU+ cells in each OB layer show a significant increase in BrdU+ cells in EPL of mutant mice compared to controls, however, no other significant increase was observed in the remaining layers of the OB. **(F, G)** Immunofluorescence staining against dorsal V-SVZ derived interneuron cell marker, Calr+, shows no visual difference in total number of Calr+ cells in the OB as a whole or in individual layers of the OB between mutants and controls. **(H)** Quantification of Calr+ cells in the OB confirms there’s no significant difference in individual layers of the OB or as a whole. **(I, J)** Immunofluorescence staining against ventral V-SVZ derived interneuron cell marker, Calb+, shows no visual difference in total number of Calb+ cells in the OB as a whole or in individual layers of the OB between mutants and controls. **(K)** Quantification of Calb+ cells in the OB confirms there’s no significant difference in individual layers of the OB or as a whole. **p-value ≤ 0.03; GL, glomerular layer; EPL, external plexiform layer; MCL, mitral cell layer; IPL, internal plexiform layer; GCL, granule cell layer

### Normal migration and maturation of olfactory bulb (OB) interneurons in the P28 *hGFAP*^*cre/+*^*;Sufu*^*fl/fl*^ mice

The dramatic changes in Gsx2+ and Tbr2+ TACs, as well as the accumulation of cells in the V-SVZ, led us to investigate whether TACs were able to transition into immature neurons, migrate through the RMS, and differentiate into interneurons that integrate into the OB circuitry. We immunostained for Doublecortin (Dcx) to examine distribution of immature nueorns and found that Dcx-expressing (Dcx+) cells were abundant along the dorsal V-SVZ and the RMS indicating that TACs were able to differentiate into immature neurons (**Figure 7A**). To confirm that production of interneurons by NSCs and TACs in the dorsal V-SVZ of *hGFAP*^*cre/+*^*;Sufu*^*fl/fl*^ mice, we examined the progenies of dividing cells in the P7 control and mutant V-SVZ. We did this by treating neonatal pups with 3 pulses of BrdU every 12 hours beginning at P6 to efficiently label proliferating precursor cells and their localization and identity within the OB (**Figure 7B**). Results from these experiments showed that BrdU-labeled V-SVZ cells successfully migrated and integrated into various OB layers by P28 (**Figure 7C-7D**). We found that the total number of BrdU+ cells in the OB of control and mutant mice did not significantly differ (**Figure 7E**, 0.08992 ± 0.01195 cells per 100 μm^2^ for n=3 controls and 0.08334 ± 0.004048 cells per 100 μm^2^ for n=3 mutants; p-value=0.6297), indicating that immature neurons originating from the P7 *hGFAP*^*cre/+*^*;Sufu*^*fl/fl*^ V-SVZ were able to migrate into the OB. Quantification of BrdU+ cells in each OB layer revealed a significant increase in BrdU+ cells in the external plexiform layer (EPL) of mutant mice compared to controls (**Figure 7E**, 0.02938 ± 0.003135 cells per 100 μm^2^ for n=3 controls and 0.04247 ± 0.001264 cells per 100 =m2 for n=3 mutants; p-value=0.0180). However, no significant changes were observed in the number of BrdU+ cells of the granule cell layer (GCL) (0.1309 ± 0.01715 cells per 100 μm^2^ for n=3 controls and 0.1098 ± 0.01342 cells per 100 μm^2^ for n=3 mutants; p-value=0.3871), internal plexiform layer (IPL) (0.06176 ± 0.01272 cells per 100 μm^2^ for n=3 controls and 0.07677 ± 0.00666 cells per 100 μm^2^ for n=3 mutants; p-value=0.3550), or mitral cell layer (MCL) (0.1176 ± 0.02794 cells per 100 μm^2^ for n=3 controls and 0.1147 ± 0.01621 cells per 100 μm^2^ for n=3 mutants; p-value=0.9313), between controls and mutants. These findings indicate that an equal number of neurogenic precursors from the dorsal V-SVZ were able to migrate into the OB despite the drastic changes in the number of neurogenic precursors.

To examine whether specific interneuron populations were affected by defects in the dorsal V-SVZ of neonatal *hGFAP*^*cre/+*^*;Sufu*^*fl/fl*^ mice, we conducted immunostaining for Calretinin, which labels interneurons known to originate from Gsx2+ cells of the dorsal V-SVZ. As shown in **Figure 7F** and 7G, Calretinin-expressing (Calr+) cells were present across all OB layers of control and mutant P28 OB. Indeed, we did not find any difference in the total number of Calr+ cells in the OB as a whole (0.07293 ± 0.009564 cells per 100 μm^2^ for n=3 controls and 0.05808 ± 0.002665 cells per 100 μm^2^ for n=3 mutants; p-value=0.2090) or in each OB layer between controls and mutants (**Figure 7H**). Additionally, we analyzed Calbindin-expressing (Calb+) interneurons, which are typically generated by neuronal precursors of the ventral V-SVZ. We found no obvious difference in the distribution of Calb+ neurons across all OB layers (**Figure 7I-7J**). We also did not find any significant differences in the total number of Calb+ neurons between control and mutant OB (**Figure 7K**, 0.3904 ± 0.08594 cells per 100 μm^2^ for n=3 controls and 0.3273 ± 0.08067 cells per 100 μm^2^ for n=3 mutants; p-value=0.5966), nor in any specific OB layer. Taken together, these findings showed that the dramatic expansion of the dorsal V-SVZ in the neonatal *hGFAP*^*cre/+*^*;Sufu*^*fl/fl*^ mice, did not inhibit the generation of properly specified and functional Type A cells. Indeed, as with control mice, these cells retained the capability to migrate and mature into specific interneuron subtypes in the OB of *hGFAP*^*cre/+*^*;Sufu*^*fl/fl*^ mice.

### Increased cell death in the P7 *hGFAP*^*cre/+*^*;Sufu*^*fl/fl*^ mice dorsal V-SVZ

The lack of any significant changes in OB interneurons prompted us to further examine the fate of NSCs and TACs in the dorsal V-SVZ of *hGFAP*^*cre/+*^*;Sufu*^*fl/fl*^ mice. We previously observed similar expansion of specific cortical progenitors in the embyronic neocortex of conditional Sufu knockouts and found that many were unable to survive (Yabut *et al.*, 2016). Therefore, we investigated whether cells in the dorsal V-SVZ of the neonatal *hGFAP*^*cre/+*^*;Sufu*^*fl/fl*^ mice were similarly unstable and became apoptotic. We conducted immunostaining against the cell death marker, Cleaved Caspase 3, and observed many dying cells (Cl-Casp3+) along the dorsal V-SVZ of mutant mice whereas Cl-Casp3+ cells in the dorsal V-SVZ of control mice were less frequent (**Figure 8A-8D**). Indeed, quantification of Cl-Casp3+ cells reflected these observations and showed a significant increase in apoptotic cells in the dorsal V-SVZ of P7 *hGFAP*^*cre/+*^*;Sufu*^*fl/fl*^ mice (**Figure 8E**; 0.184 ± 0.08815 cells per 100 μm^2^ for n=3 controls and 0.5282 ± 0.0672 cells per 100 μm^2^ for n=3 mutants; p-value=0.0360). Altogether, these findings suggest that despite the massive expansion of precursor cells in the dorsal V-SVZ, many of these cells failed to survive and differentiate into mature cell types.

**Figure 8.**
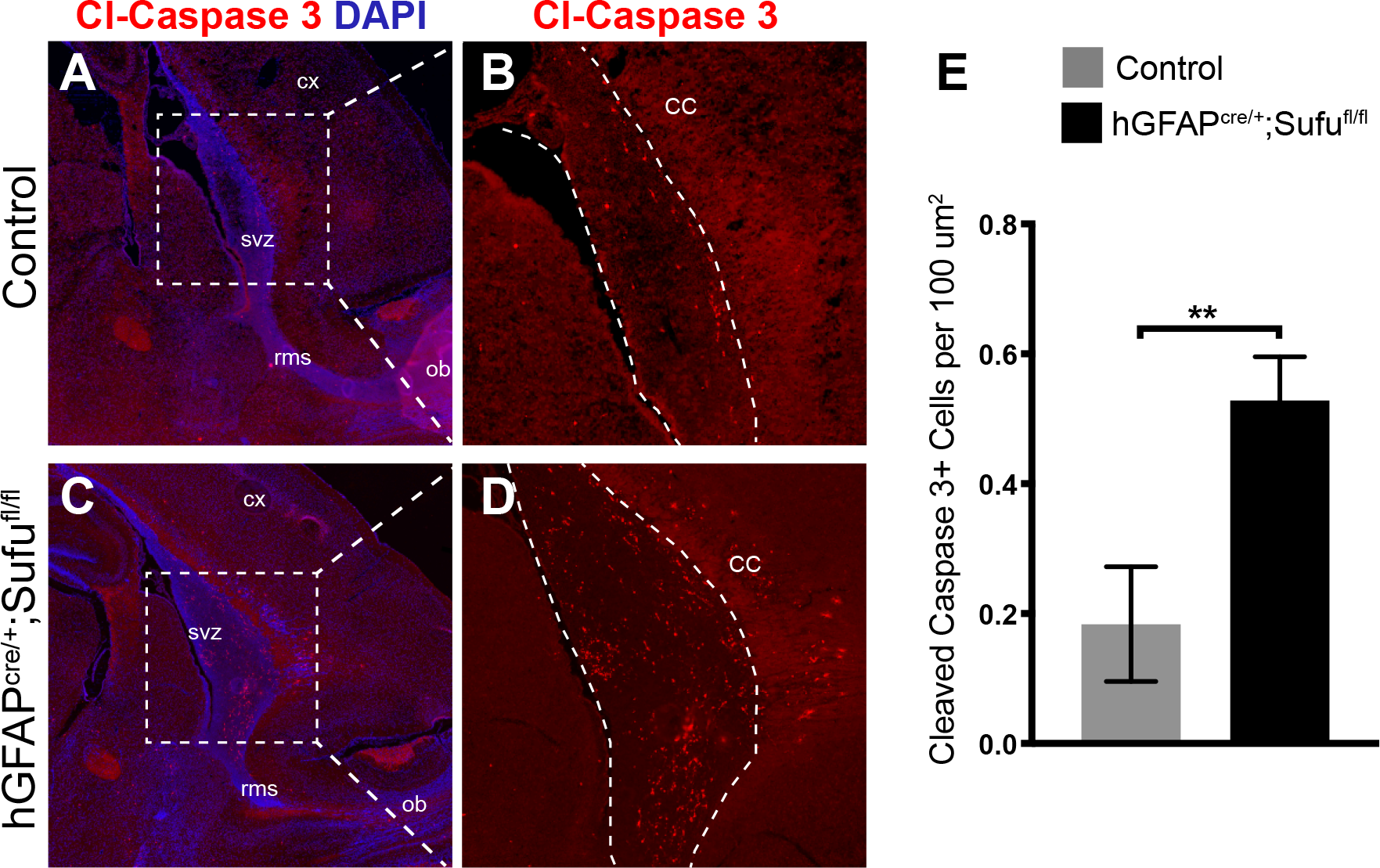
Increase in cell death along the SVZ of the P7 *hGFAP*^*cre/+*^*;Sufu*^*fl/fl*^ mice. **(A-D)** Immunofluorescence staining against cell death marker, Cleaved Caspase 3 (Cl-Casp3+), shows an observable difference in the number of Cl-Casp3+ in the dorsal V-SVZ of P7 *hGFAP*^*cre/+*^*;Sufu*^*fl/fl*^ mice compared to controls. **(D)** Quantification of Cl-Casp3+ cells in the V-SVZ demonstrated a significant increase in P7 *hGFAP*^*cre/+*^*;Sufu*^*fl/fl*^ mice compared to controls. **p-value ≤ 0.03; V-SVZ, ventricular-subventricular zone; RMS, rostral migratory stream; OB, olfactory bulb; CX, cortex

## DISCUSSION

The postnatal V-SVZ is composed of multiple neuronal precursor populations that sustain lifelong neurogenesis in rodents. Our study provides insights into the molecular mechanisms involved in the formation of a molecularly distinct neurogenic domain, the dorsal V-SVZ. We showed that the cytoplasmic adaptor protein, Sufu, plays important roles in controlling precursor number and viability. Genetic ablation of Sufu in RG cells at late embryonic stages caused a dramatic expansion of the dorsal V-SVZ, but not the ventral V-SVZ. This expansion is due to the uncontrolled proliferation and organization of Sox2+ Type B cells, resulting to the deregulated production of TACs via Gli3-dependent mechanisms, and independent of Shh signaling activity. Our novel findings establish a crucial role for Sufu in maintaining precursor populations in the neonatal dorsal V-SVZ.

Our results emphasize that differential regulatory mechanisms are employed to establish and maintain molecularly distinct dorsal and ventral V-SVZ regions (Merkle, Mirzadeh and Alvarez-Buylla, 2007). We found that loss of Sufu at late embryonic stages did not disrupt the formation of the V-SVZ. Mice lacking Sufu formed anatomically distinct dorsal and ventral V-SVZ domains capable of generating predicted subpopulations of interneuron subtypes in the OB. This indicates that progenitor specification, as determined by their localization along the dorsoventral V-SVZ axis, was not severely disrupted. However, the dorsal V-SVZ was expanded in mice lacking Sufu, as a result of the persistent proliferation of Sox2+ Type B1 cells. These findings implied that loss of Sufu at late embryonic stages maintained NSCs in a highly proliferative state at neonatal stages, producing abnormally large numbers of Gsx2+ TACs specifically in the dorsal V-SVZ. Thus, Sufu must play a role in modulating the cell cycle progression of NSCs in the dorsal V-SVZ to ensure a timely production of specific NSC lineages.

Loss of Sufu differentially affected specific neural progenitor populations based on their embryonic origins. Tbr2+ neural progenitors that typically originate from the embryonic cortical progenitors were significantly reduced whereas Gsx2+ neural progenitors that originate from the embryonic ganglionic eminence (Stenman, Toresson and Campbell, 2003; Brill *et al.*, 2009) were significantly increased. We previously reported that loss of Sufu results in the increase in proliferation of cortical progenitors (resulting in an increase in superficial layer projection neurons in the neocortex) and oligodendrogenesis in the E16.5 neocortex of hGFAP-Cre;Sufu-fl mice (Yabut *et al.*, 2016; Winkler *et al.*, 2018). Thus, two possibilities could explain the reduction in Tbr2+ progenitors: 1) cortical progenitors that generate V-SVZ NSCs were re-specified towards the gliogenic lineage, and/or 2) exhaustion of the cortical RG pool has occurred. This would prevent RG cells to generate Type B1 cells in the dorsal V-SVZ. Lineage tracing of embryonic cortical progenitors will determine if proportions of RG cells in the E16.5 neocortex, a time at which Type B1 cells are thought to be specified (Fuentealba *et al.*, 2015; Furutachi *et al.*, 2015), failed to generate Type B1 cells in the neonatal V-SVZ. Nevertheless, these findings indicate that in addition to previously identified roles of Sufu in corticogenesis, Sufu also functions to ensure the proper production of dorsal V-SVZ NSCs that generate specific subtypes of TACs. The consequence of this abnormality is unclear since we did not find any major changes in the generation of specific OB interneuron subtypes. However, given the molecular heterogeneity of OB interneurons, we cannot exclude the possibility of physiological abnormalities that could arise from abnormal maturation and integration of specific subsets of OB interneurons.

Sufu antagonizes Shh signaling by mediating the proteolytic processing of Gli transcription factors to either inhibit activator function or promote the formation of transcriptional repressor forms. In the developing forebrain, we have previously shown that Sufu acts to regulate the stability and processing of Gli2 and Gli3 proteins at early stages of corticogenesis that resulting in an increase in Gli2R and Gli3R levels, while it functions to promote the generation of Gli3R alone at later stages (Yabut *et al.*, 2015). Here, we found that loss of Sufu affected Gli transcription in dorsal V-SVZ cells. We found that Gli1 and Gli3 mRNA levels were significantly reduced in the *hGFAP*^*cre/+*^*;Sufu*^*fl/fl*^ mice V-SVZ. Gli3 is typically highly expressed in the dorsal V-SVZ to exert its repressor function during the establishment of the V-SVZ (Petrova, Garcia and Joyner, 2013; Wang *et al.*, 2014). These findings showed that in the absence of Sufu, Gli3 transcription is not efficiently maintained in the neonatal V-SVZ. Loss of Sufu may have resulted in the repression of Gli3 expression by deregulation of yet unidentified transcription factors that typically promote Gli3 transcription, or indirectly caused by defects in Gli3 protein processing triggering other transcriptional repressors to inhibit Gli3 expression. Elucidating the molecular steps by which Sufu alters Gli3 expression and activity in dorsal V-SVZ NSCs could provide novel insights on the diverse mechanisms utilized by Sufu to control Gli3 function.

Deletion of Sufu and the subsequent reduction in Gli3 transcription in the neonatal *hGFAP*^*cre/+*^*;Sufu*^*fl/fl*^ mice resulted in defects that phenocopy Gli3R-cKO mice (Wang *et al.*, 2014). Similar to our findings, ectopic activation of Shh signaling did not occur in Gli3R-cKO V-SVZ at neonatal stages. This could be because responsiveness to Shh signals by neonatal dorsal V-SVZ cells, such as in NSCs and TACs, do not occur until after P7 (Ahn and Joyner, 2005; Palma *et al.*, 2005; Wang *et al.*, 2014). Supporting this, conditional deletion of Smo in V-SVZ NSCs at P0, using the mGFAP-Cre driver, does not cause any obvious proliferation defects in the V-SVZ until after P15 (Petrova, Garcia and Joyner, 2013). This would indicate that the Sufu-Gli3 regulatory axis alone is critical in the control of progenitor populations in the dorsal V-SVZ niche, independent of its canonical roles as regulators of Shh signaling activity. Indeed, previous studies have shown that Gli3R functions to regulate gp130/STAT3 signaling in NSCs at early neonatal stages for proper establishment of the V-SVZ niche (Wang *et al.*, 2014).

In summary, our studies identified multiple roles for Sufu in establishing appropriate cell number and identity in the neonatal dorsal V-SVZ. We found that Sufu maintains neurogenic precursor populations in the dorsal V-SVZ via regulation of Gli3. These findings underscore the importance of Sufu as a key regulator of stem/progenitor populations not only in the developing embryonic forebrain but also in establishing postnatal neurogenic niches such as the V-SVZ. These results have potential implications in how neural stem/progenitor populations are established and sustained in the postnatal neurogenic niche, how defects in proliferation could predispose these cells to a number of neurological diseases and malignancies, and provide insights on potential molecular strategies that can be utilized for regenerative therapies.

## AUTHOR CONTRIBUTIONS

HG performed experiments, analyzed the data, and wrote the manuscript. JGC, HN, DA performed experiments and analyzed the data. ORY and SJP conceived of and performed experiments, analyzed the data, and wrote the manuscript.

## ACKNOWLEDGMENTS

We would like to thank members of the Pleasure Lab for helpful discussions, Dr. Kenneth Campbell for the Gsx2 antibody, and DeLaine Larsen and Kari Harrington at the University of California San Francisco Nikon Imaging Center for assistance with imaging.

